# Population genetic analysis of Indian SARS-CoV-2 isolates reveals a unique phylogenetic cluster

**DOI:** 10.1101/2020.07.19.197129

**Authors:** Dhruv Das, V.S.S.N.R Akkipeddi

## Abstract

The SARS-CoV-2 pandemic originated from Wuhan, China in December 2019 raised an alarming situation all over the globe. Sequencing of this novel virus provides an opportunity to evaluate the genetic polymorphism present in the viral population. Herein, we analysed 173 sequences isolated from Indian patients and performed SNP linkage, clustering and phylogenetic analysis to understand the local genetic diversity. We found that the SNP linkages that lead to the identification of some global clades, do not hold true for the local clade classification. In addition to the unique cluster, established by another Indian study, we identified a new cluster (I-20D) that encompasses 28% of the analysed sequences. This cluster is defined by two linked variations – C22444T and C28854T. A detailed study of such polymorphisms can be useful for drug and vaccine development.

## Introduction

The emergence and global spread of the novel Severe Acute Respiratory Syndrome Coronavirus 2 (SARS-CoV-2) causing coronavirus disease 2019 (COVID-19) is the major point of concern for all affected countries. The origin of SARS-CoV-2 can be traced back to late 2019 in Wuhan, China^1^. Since then SARS-CoV-2 has spread across the world and become a global health emergency on March, 2020 when World Health Organisation (WHO) declared COVID-19 as global pandemic^2^. The worldwide alarming situation can be foreseen with more than 9,500,000 people infected and over 489,000 deaths reported globally (as of June 26^th^ 2020). After its entry into human host, the continuous divergence of the virus generated at least 5 major clades globally till June 2020^3^. Genetic analysis of the virus population can provide information about the extent of molecular divergence and spread of SARS-CoV-2 in the human population.

Before December 2019, six human coronaviruses were known with different severity causing pulmonary and gut infections. Four out of six, HCoV-229E, HCoV-NL63, HCoV-OC43, HCoV-HKU1 were common coronaviruses that are infecting worldwide population with low pathogenicity to humans^4,5^. The remaining two human coronaviruses, Severe Acute Respiratory Syndrome Coronavirus (SARS-CoV) and Middle East Respiratory Syndrome Coronavirus (MERS-CoV) are the two highly pathogenic coronaviruses. The novel seventh Coronavirus (SARS-CoV-2) is a close relative of the SARS-CoV species as they share high homology (~80%) and recognize angiotensin-converting enzyme 2 (ACE2) to initiate infection cycle^5^. SARS-CoV-2 is a positive-sense ssRNA (single stranded RNA) virus with approximately 30kb genome size.

Similar to SARS-CoV and MERS-CoV, bats serve as reservoir hosts for progenitor of SARS-CoV-2^6^. SARS-CoV-2 shares the highest sequence homology (96.3%) with bat coronavirus (RaTG13) isolated from a bat in Yunnan^7^. The Receptor Binding Domain (RBD) of SARS-CoV-2’s surface glycoprotein shares (86.64%) nucleotide sequence identity with pangolin CoV’s RBD as compared to the bat virus, RaTG13 (86.19%)^8^. A recent study highlights a recombination event between bat CoV and pangolin CoV, before its introduction in humans^9^. Strong purifying selection pressure and recombination shapes the origin and evolution of the novel SARS-CoV-2^6,9^. After its human entry the molecular divergence of SARS-CoV-2 is continuing and it is important to track this molecular evolution to understand the epidemiology of the disease and to effectively design drugs and vaccines for a target population.

In an earlier study^10^, 103 SARS-CoV-2 genomes were segregated in two lineages ‘L’ and ‘S’ based on two linked mutations. The expansion of the viral genome sequences revealed large number of polymorphism in the population and their geo-distribution^11^. Nextstrain defines five major global clades that can classify the overall global SARS-CoV-2 genome sequences^3^. The geo-distribution of the mutations and therefore clades^11^ across countries can have severe impact on the local evolution of the virus. In a previous study^12^, combination of four genetic variants were used to define a unique clade predominant in India.

The publicly available whole genome sequences of SARS-CoV-2 isolates from different countries can be utilized for understanding the evolution of the virus as of function of time and geography^11^. The information can be used to characterize the novel viral mutations in a population that can be further exploited for designing better diagnostic tests, vaccines and antiviral drugs. In the present study, we characterized the genetic variants in 173 SARS-CoV-2 genomes isolated from the Indian population, predominantly from Ahmedabad and Gujarat. We analysed the genetic variants defining the phylogenetic clusters and established a new clade dominant in the Indian population that was not reported earlier.

## Materials and Methods

### Data collection and sequence analysis of SARS-CoV-2 isolated from Indian Population

173 FASTA sequence files of SARS-CoV-2 were retrieved from NCBI^13^ (update: 05-06-2020) database. Only complete genomes originated from Indian population were used in this study. Reference sequence and annotations of SARS-CoV-2 (NC_045512) were retrieved from GenBank, NCBI^13^ and the reference sequence was used as parent sequence for all other sequences. The sequences were divided into two sets: initial set (update: till 17-05-2020) and final set (update: till 05-06-2020). The first set of 60 sequences (59 Indian originated sequences and 1 reference sequence) were used for initial analysis of genetic diversity, SNP linkages, clustering and phylogenetic analysis. And the final set of 174 sequences was used to test the results concluded from the analysis of initial set.

Multiple Sequence Alignment (MSA) of initial set sequences was performed using MUSCLE algorithm in MEGA^14^. Further, MEGA^14^ was used to evaluate the mutation type (Synonymous or Non-synonymous) and frequencies at every single site in aligned sequences of whole genome as well as concatenated CDS (coding regions). Mutations occurring 5 or more than 5 times were considered as recurrent mutations. The protein and conserved domain information were retrieved from NCBI^15^.

### Linkage Disequilibrium and linkage-based sequence clustering

The aligned file of all retrieved sequences of initial set was used to generate a HapMap file with sixteen selected mutations containing information regarding sequence, sequence position and allele frequency of both minor and major. TASSEL version 5.2.61^16^ was used to generate linkage plot between the selected SNPs using HapMap file. Correlation analysis between mutations, sequences and geographical locations was performed using ClustVis^17^ tool. A heat map was generated using the following settings: unit variance scaling was applied to rows, both rows and columns were clustered using correlation distance and average method, and tightest cluster first was used for tree ordering of both rows and columns. Clusters were defined based on the linked SNPs in the sequence.

### Phylogenetic analysis and global distribution of clades

Phylogenetic analysis between SARS-CoV-2 sequences used in this study was performed using PhyML^18^ with Smart Model Selection^19^ with tree searching using Nearest Neighbour Interchange (NNI) and 100 bootstrap replicates. GTR+I substitution model was selected as best model using BIC (Bayesian Information Criterion) selection criteria.

Nextstrain nomenclature^3^ for clade identification was adopted and used to classify the sequences in this study. For defining a new clade, at least 2 mutations should be away from its parent major clade.

For global analysis, sequences were retrieved from all the countries who had deposited in NCBI till June 5^th^ 2020. Then Multiple sequence alignment (MSA) was performed and frequency of the occurrence of the two mutations simultaneously was assessed.

### Functional Characterization of amino acid change

The N-gene variant, C194T, was evaluated for the functional characterization using the Protein Variant Effect Analyser (PROVEAN) and PolyPhen-2. In PROVEANv1.1^20^, the predefined threshold score was used, i.e. −2.5. This cut off score corresponds to 80.4% sensitivity and 78.6% specificity. Polyphen-2 uses Naïve Bayes classifier for predicting functional significance of the allele replacement^21^. Default parameters has been used for calculating the Polyphen-2 score.

## Results

### SARS-CoV-2 genome sequences, their origin and date of collection

The viral genome isolates of Indian origin (n=173) along with the reference genome sequence (Wuhan-Hu-1, NC_045512.2) were collected over the period of five and half months from December 24, 2019 to June 5, 2020; and most of them (94.8%) were isolated after mid-April (**Fig: 1a**). The most represented origin of isolated viruses was Ahmedabad, with 43.3% (75/173) share of all sequences (**Fig: 1b**). Out of these 173 Indian origin SARS-CoV-2 sequences, 59 SARS-CoV-2 complete genomes (update: till 17-05-2020; initial set) were used for initial evaluation of genetic diversity, SNP linkages and phylogenetic analysis. Rest of the sequences along with the initial set (n = 173) were later used to validate the results obtained from the analysis of initial sequence set.

**Fig: 1a.**
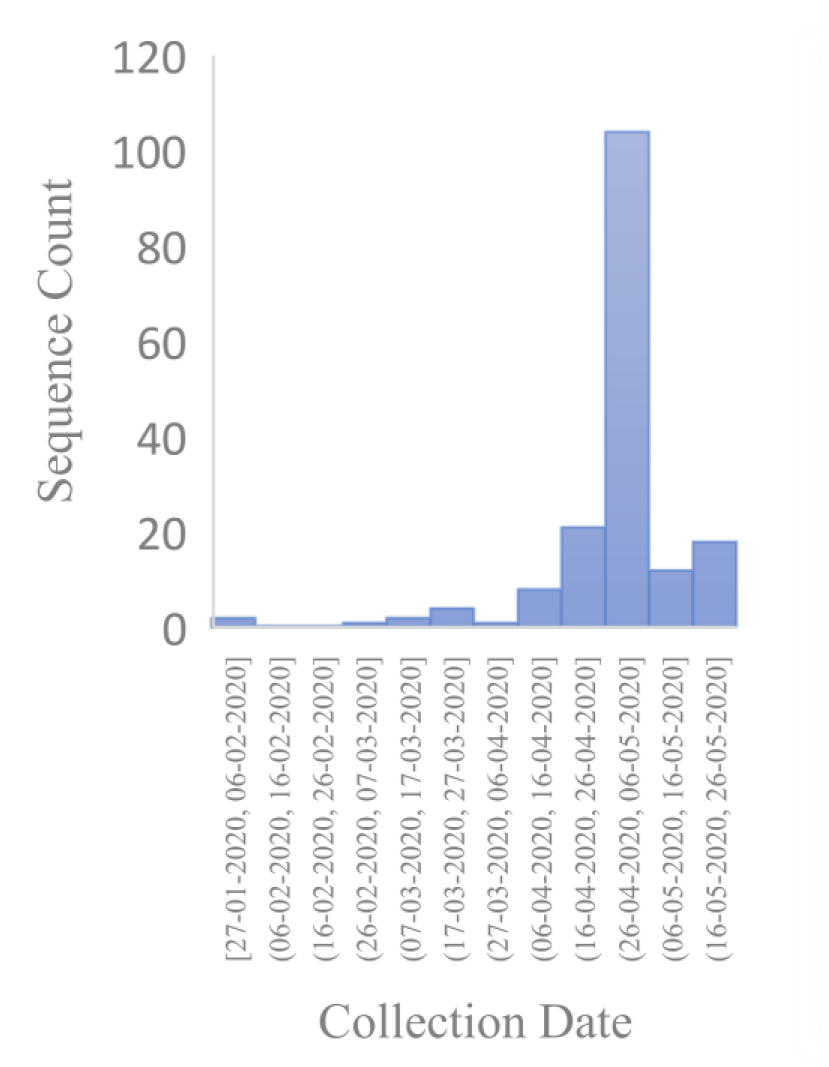
The histogram represents the collection date of viral genomes in Indian population over a period of 5 and half months from December 24, 2019 to June 5, 2020

**Fig: 1b.**
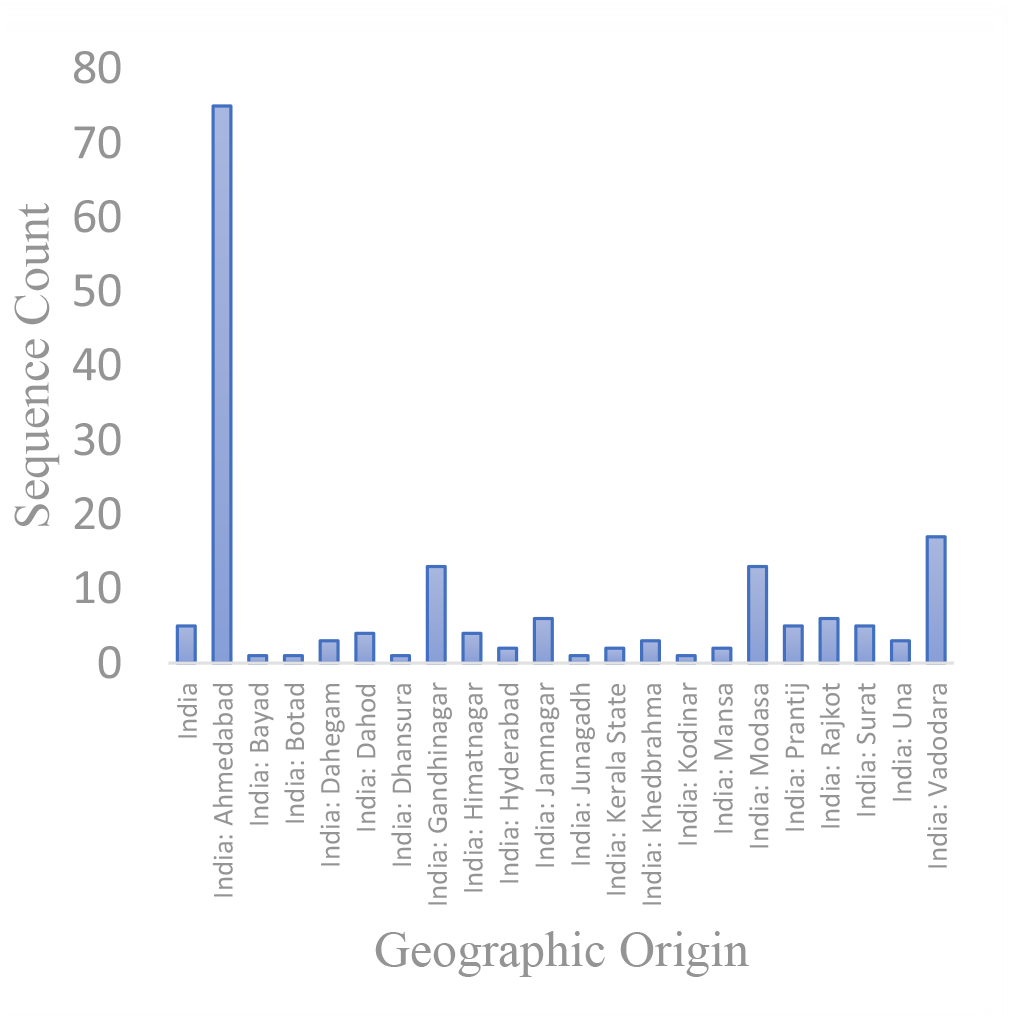
Geographic origin of the sequences used in the study.

### Genetic diversity in the Indian SARS-CoV-2 sequences

A total of 136 variations were identified in the initial sequence set studied as compared to the reference sequence Wuhan-Hu-1/2019 (NC_045512.2), Additionally, two unrelated 3-nucleotide deletions were observed in two different sequences. There were 91 singleton mutations present, harbouring change only in one sequence. There were four sites, one in the 5’ (1-265) and three in the 3’ (29675..29903) UTR regions where mutations were observed. All mutations present in 3’ UTR region were singleton. However, a C to T mutation at position 241 (compared to the reference genome) in the 5’-UTR was found in about 86% of the 173 Indian sequences.

In concatenated CDS region of all genes, total 131 variations were found, including 82 (62.6%) non-synonymous and 49 (37.4%) synonymous mutations (**Fig: 2a**). Out of them, 88 (67.2%) were singleton that includes 55 (62.5%) non-synonymous and 33 (37.5%) synonymous mutations (**Fig: 2a**). For closely related isolates, relationship between selection and dN/dS values is not monotonic. The high occurrence of non-synonymous mutations (**Fig: 2b**) at selected sites might suggest a rapid sweep in the population and strong selection^22^. Therefore, predicting selection pressure from dN/dS can be misleading^22^.

**Fig: 2a.**
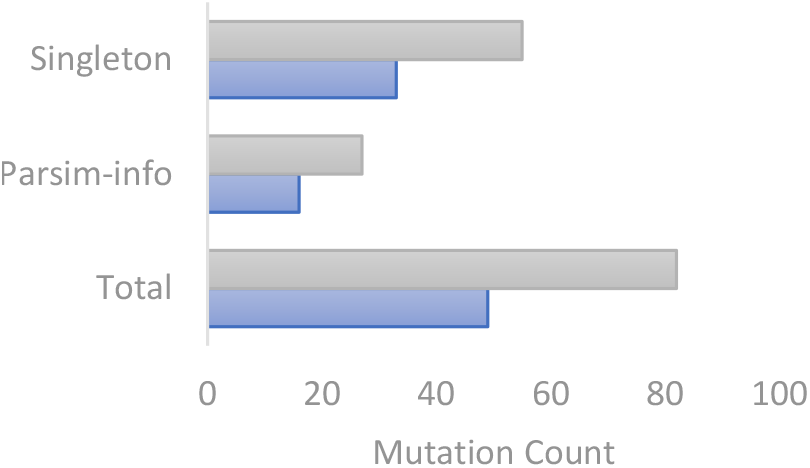
Mutation spectra summary in the initial sequence set. Singleton refers to the mutations occurring once in the sequence population. Mutations occurring more than once are referred as parsim-info.

**Fig: 2b.**
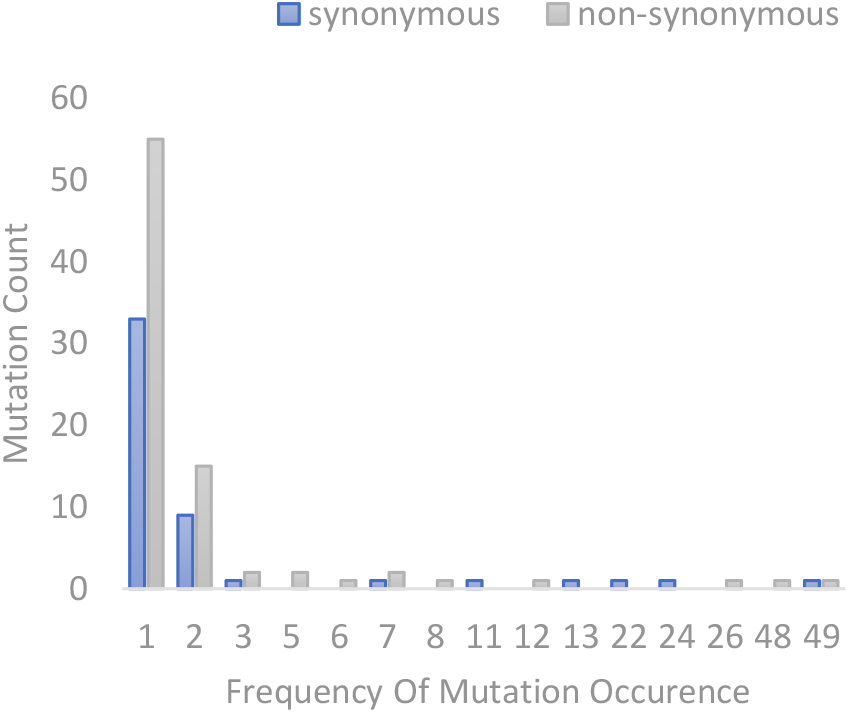
Synonymous v/s non-synonymous mutation distribution in the sequences of initial set.

The analysis of the frequencies of mutated alleles in the initial set revealed the presence of 16 frequent mutations with minor allele frequency of f > 5% (the frequency that is considered significant^23^), i.e. mutations present in more than three SARS-CoV-2 genomes were considered for this analysis. The frequencies of these 16 selected recurrent mutations for initial as well as final set are given in **Table: 1**.

**Table: 1.** General details about the selected recurrent mutations and comparison between frequency of recurrent mutation occurrence in initial and final sequence set.

| S. No. | Genomic Location<br>(Base Change) | ORF / Protein<br>(Codon Change) | Change in<br>Amino Acid | Frequency w.r.t<br>Reference<br>Sequence<br>(n=59) | Frequency w.r.t<br>Reference<br>Sequence<br>(n=173) |
| --- | --- | --- | --- | --- | --- |
| 1 | 2836 (C>T) | ORF1ab / nsp3<br>(TGC857TGT /<br>TGC39TGT) | Synonymous | 18.6% | 22% |
| 2 | 3037 (C>T) | ORF1ab / nsp3<br>(TTC924TTT /<br>TTC106TTT) | Synonymous | 83% | 86.7% |
| 3 | 4866 (G>T) | ORF1ab / nsp3<br>(AGT1534ATT /<br>AGT716ATT) | S1534I/S716I | 8.8% | 3.5% |
| 4 | 6312 (C>A) | ORF1ab / nsp3<br>(ACA2016AAA /<br>ACA1198AAA) | T2016K/T1198K | 11.9% | 8.7% |
| 5 | 11083 (G>T) | ORF1ab / nsp6<br>(TTG3606TTT /<br>TTG37TTT) | L3606F/L37F | 10.2% | 11% |
| 6 | 13730 (C>T) | ORF1ab/RNA-dependent<br>RNA polymerase<br>(GCT4489GTT /<br>GCT97GTT) | A4489V/A97V | 11.9% | 8.1% |
| 7 | 14408 (C>T) | ORF1ab/RNA-dependent<br>RNA polymerase<br>(CCT4715CTT /<br>CCT323CTT) | P4715L/P323L | 83% | 85% |
| 8 | 14425 (C>A) | ORF1ab/RNA-dependent<br>RNA polymerase<br>(CTA4721ATA /<br>CTA329ATA) | L4721I/L329I | 8.8% | 3.5% |
| 9 | 18877 (C>T) | ORF1ab polypeptide /3'-<br>to-5' exonuclease<br>(CTA6205TTA) | Synonymous | 40.7% | 52% |
| 10 | 22444 (C>T) | S-gene/surface<br>glycoprotein<br>(GAC294GAT) | Synonymous | 22% | 28.3% |
| 11 | 23403 (A>G) | S-gene/surface<br>glycoprotein<br>(GAT614GGT) | D614G | 81.3% | 86.1% |
| 12 | 23929 (C>T) | S-gene/surface<br>glycoprotein<br>(TAC789TAT) | Synonymous | 11.9% | 8.7% |
| 13 | 25563 (G>T) | ORF3a protein<br>(CAG57CAT) | Q57H | 44.1% | 53.2% |
| 14 | 26735 (C>T) | M-gene/membrane<br>glycoprotein<br>(TAC71TAT) | Synonymous | 37.3% | 49.7% |
| 15 | 28311 (C>T) | N-gene/nucleocapsid<br>phosphoprotein<br>(CCC13CTC) | P13L | 13.5% | 9.2% |
| 16 | 28854 (C>T) | N-gene/nucleocapsid<br>phosphoprotein<br>(TCA194TTA) | S194L | 20.3% | 28.3 |

As can be seen from the above table, the 10 non-synonymous mutations were distributed in five genes coding seven proteins with variable frequencies, in the genes coding for ORF1ab (Nsp3, Nsp6, RNA-dependent RNA polymerase), S-gene (spike protein), ORF3a, M-gene (membrane glycoprotein) and N-gene (nucleocapsid phosphoprotein). ORF1ab being the largest coding region harboured higher non-synonymous mutations (n = 6) than any of the other four genes (each having 1 non-synonymous mutation). In ORF1ab two non-synonymous mutations were found in Nsp3 protein, one in C-terminal SARS-Unique Domain (SUD) (S716I) and the other in nucleic acid-binding domain (T1198K). In Nsp6, a transmembrane domain containing protein, of ORF1ab there was a leucine to phenylalanine mutation (L37F). The other three non-synonymous mutations in ORF1ab were present in the RPol N-terminus of RNA-dependent RNA polymerase (A97V, P323L and L329I). In surface glycoprotein there was a frequent aspartic acid to glycine change at position 614 (D614G) outside of receptor binding domain. APA3_viroporin region of ORF3a had a frequent glutamine to histidine change at position 57 (Q57H). Whereas there were two frequent amino acid changes in nucleocapsid phosphoproteins (P13L and S194L).

### SNP linkage and Phylogenetic analysis of SARS-CoV-2 genomes

To analyse and visualize the linkage pattern among the selected sixteen recurrent mutations, a linkage disequilibrium (LD) analysis was performed using TASSEL 5.2.61^16^. We found that most of the analysed SNPs had shown linkage, with value of D’ (Fig: 3a, upper right triangle) as 1 and varying R^2^ value (**Fig: 3a**, lower left triangle) as the allele frequency is varying for different mutation pairs. Some SNPs had shown strong linkage^24^ with similar allele frequencies having R^2^ value more than 0.8 along with D’ value of 1 (**Fig: 3b, Table: 2**).

**Fig: 3a.**
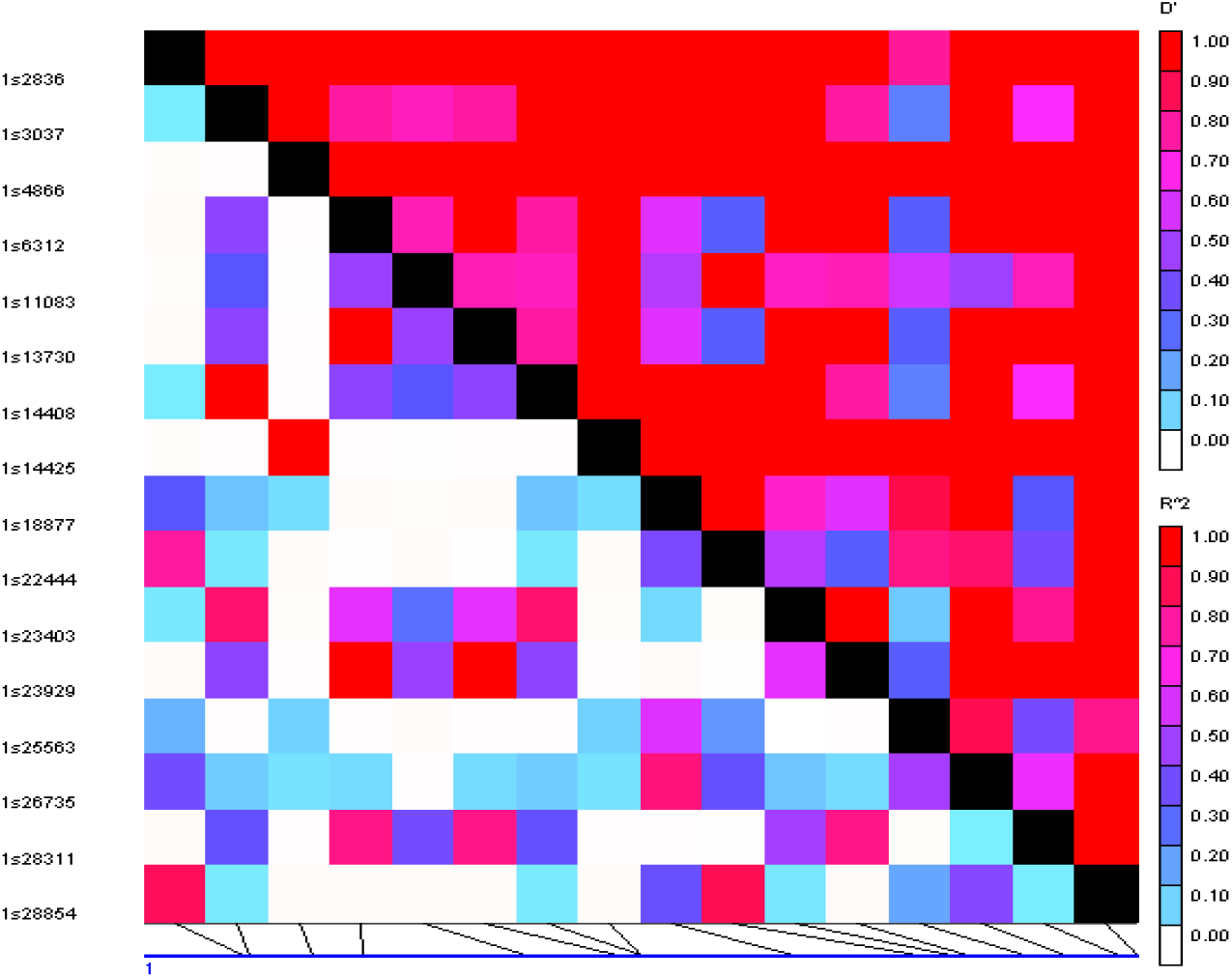
LD plot of SNP pairs among the selected 16 sites that have minor alleles in more than three sequences. The numbers on the left side of the plot shows the genomic locus of the SNPs. The upper right triangle represents the D’ value and lower left triangle represents the R^2^ value. Color scale: Red color => complete linkage; White => no linkage between two SNPs.

**Fig: 3b.**
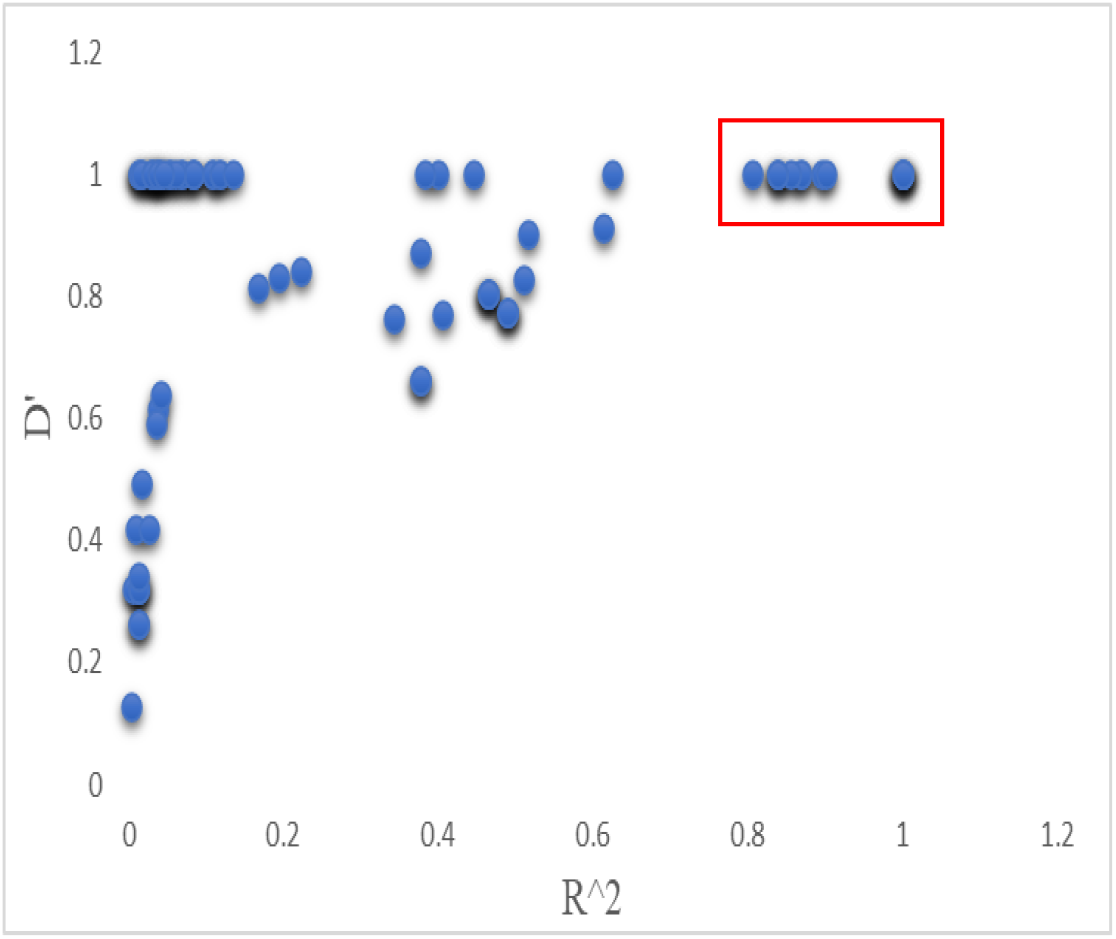
Scatter plot between the R^2^ (x-axis) and D’ (y-axis) of each SNP pair. The points in the red box had shown strong linkage with similar allele frequencies (**Table: 2**). The intensity of dark shadow behind the blue dot indicates the presence of multiple points.

**Table: 2.** R^2^ and D’ values of the SNPs marked in figure 3

| Linked SNPs position | $R^2$ | $D'$ |
| --- | --- | --- |
| 13730 - 6312 | 1 | 1 |
| 23929 - 6312 | 1 | 1 |
| 23929 - 13730 | 1 | 1 |
| 28311 - 6312 | 0.838906 | 1 |
| 28311 - 13730 | 0.838906 | 1 |
| 28311 - 23929 | 0.838906 | 1 |
| 14425 - 4866 | 1 | 1 |
| 23403 - 3037 | 0.869136 | 1 |
| 23403 - 14408 | 0.869136 | 1 |
| 14408 - 3037 | 1 | 1 |
| 22444 - 2836 | 0.805861 | 1 |
| 28854 - 2836 | 0.894841 | 1 |
| 28854 - 22444 | 0.900563 | 1 |
| 26735- 18877 | 0.855978 | 1 |

To further confirm the above finding a correlation analysis between recurrent mutations and individual sequences were performed using ClustVis^17^. As expected, we observed a clear clustering of sequences based on the linkage pattern of various mutations (**Fig: 4, Table: 2**). There were two broad clusters, one minor (A) and one major (B) cluster that divide the population in two distinct groups. The minor group was nearer to reference genome in sequence similarity and the major group can be further divided into two smaller groups (C and D). Both ‘C’ and ‘D’ encompass two further smaller clusters ‘E’ and ‘F’ respectively. Likewise, ‘A’ cluster contains a small cluster ‘G’. In our study population there was an over-representation of SARS-CoV-2 genomes isolated from Ahmedabad and most of these sequences fall in the major cluster ‘B’ indicating of a possible common origin of the sequences.

**Fig: 4.**
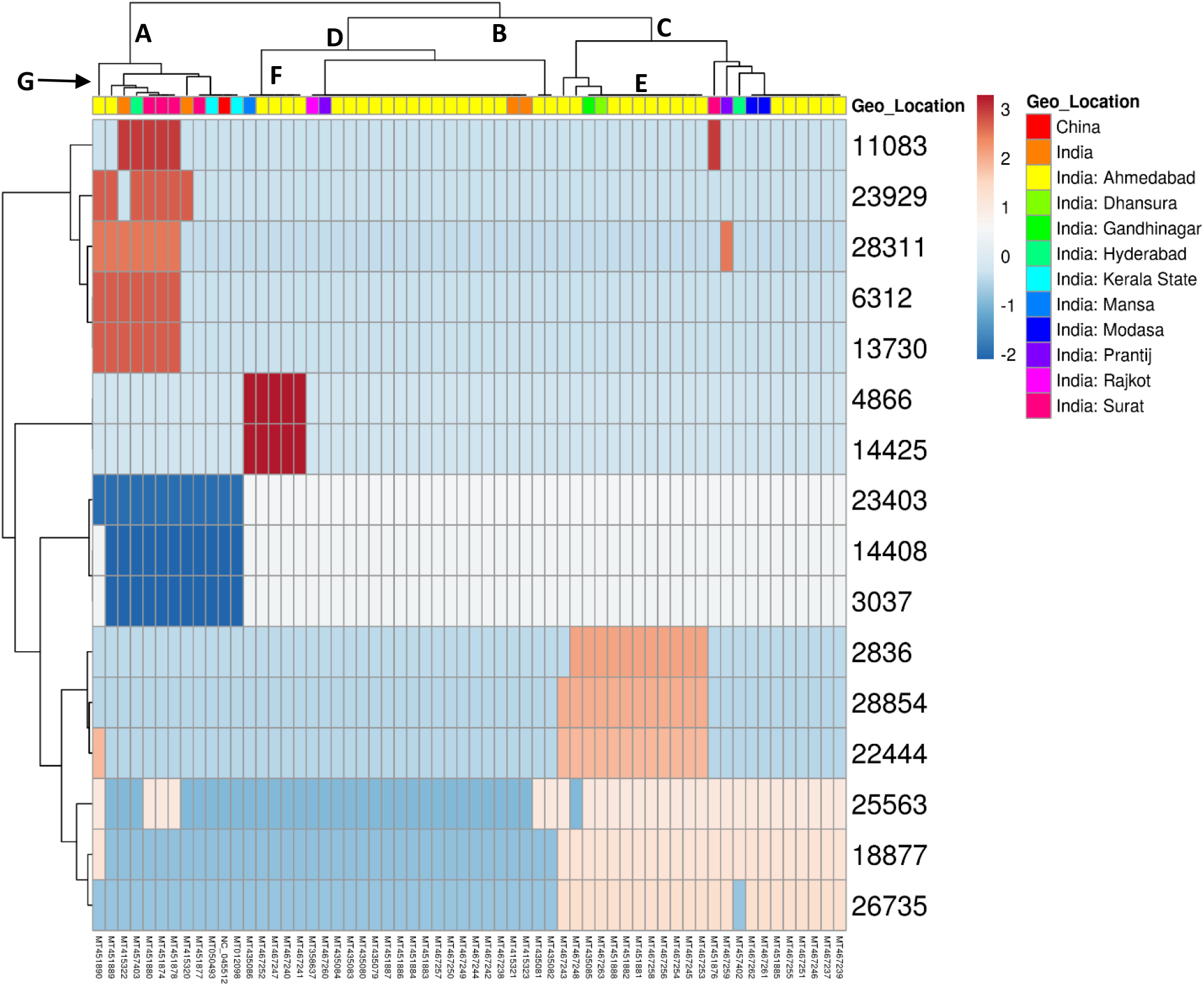
Heatmap demonstrating correlation between the selected 16 mutations and geographical origin of the analyzed sequences. The extent of correlation can be determined by color scale where blue and orange color signifies highest and lowest correlation, respectively. The colors in the extreme right horizontal bar represents the geographical origin of the sequences.

A phylogenetic tree was constructed using 60 SARS-CoV-2 whole genome sequences (including 1 reference genome) of the initial set. The phylogenetic tree revealed similar organization of clusters and in agreement with our previous results (**Fig: 5**). The lineages can be distinguished and identified based on the occurrence of the linked SNPs.

**Fig: 5.**
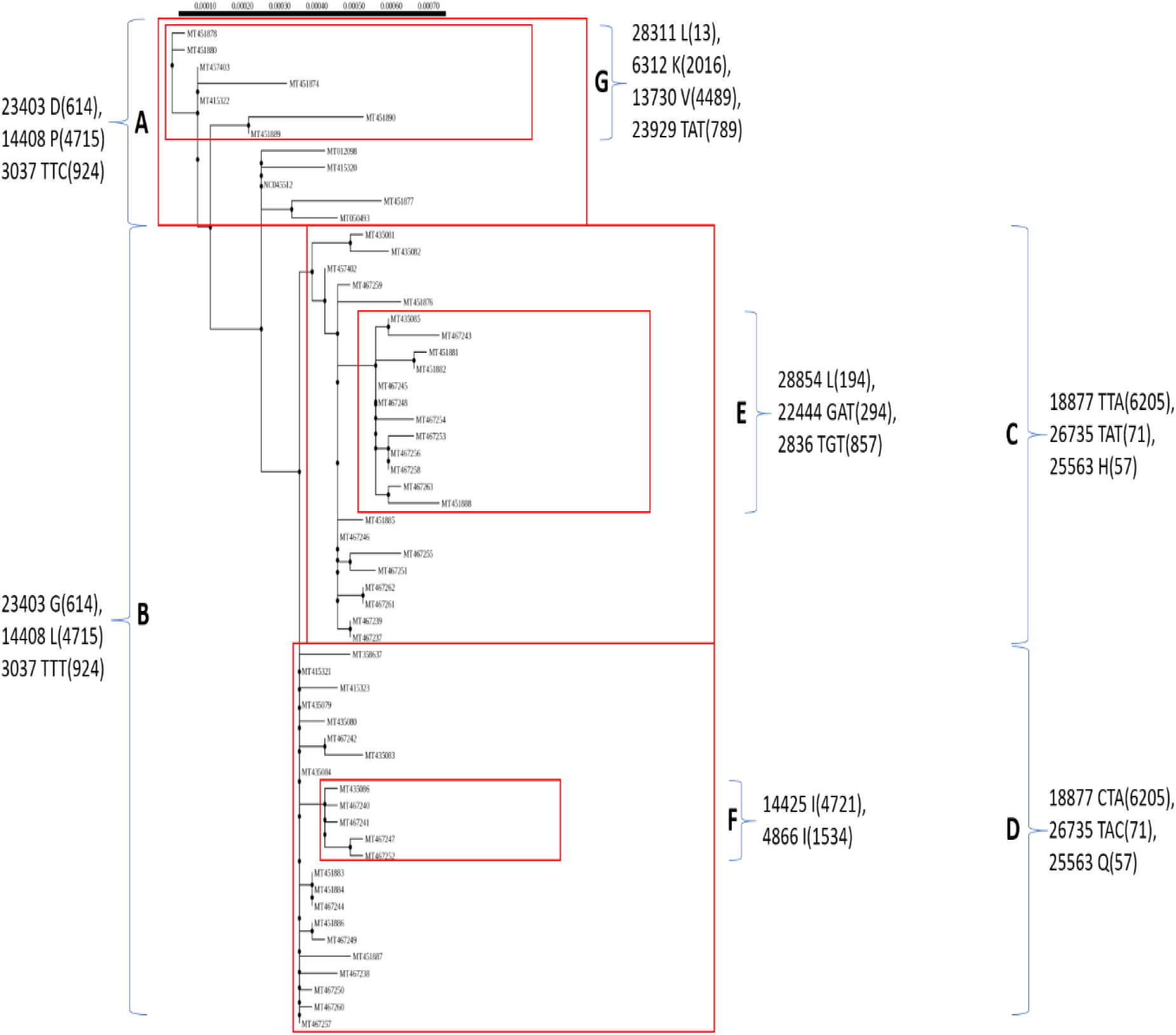
Phylogenetic analysis (100 bootstrap replications) of 60 SARS-CoV-2 genome sequences (initial set). Red boxes highlight the cluster of sequences sharing linked SNPs.

### Extending analysis to 173 sequences and comparing with the global status

When we extended our analysis by comparing all 173 sequences together, we found that most of the frequencies of selected recurrent mutations were matching with the initial sequence set (Fig: 6). Frequencies of only two non-synonymous mutations were decreased by more than five percent, indicating their loss in the population. There were some mutations which were previously present in low numbers but after extending the sequence numbers they appear with higher frequencies, suggesting either selection pressure operating on these sites or introduction of new mutations from other population. Interestingly most of these new mutations were non-synonymous in nature. Further phylogenetic analysis was performed to validate the previous clustering and phylogeny of smaller sequence set; and emergence of new clades.

**Fig: 6.**
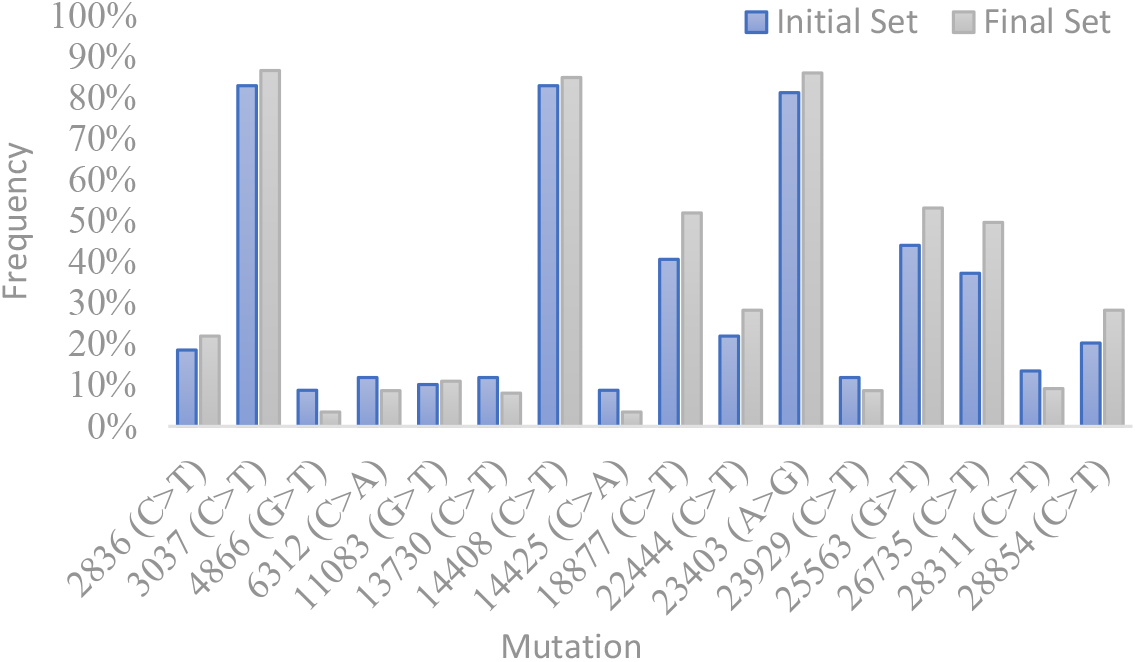
Comparison of recurrent mutation frequencies in the initial (blue) and final (grey) set.

Nextsrain^3,25^ identified five global major clades, namely 19A, 19B, 20A, 20B and 20C till June 2020. Each of these clades was defined by the presence of at least two mutations. 19A is the root clade related mostly to the reference sequence. 19B and 20A both emerged from 19A and both have two linked mutations defining their major clades, T28144C and C8782T for 19B; and C14408T and A23403G for 20A. 20B and 20C emerged from 20A defined by their linked genetic variants. G28881A, G28882A and G28883C defines 20B whereas C1059T and G25563T defines 20C.

On analysing the sequences, we found 19B, a cluster designated as a major cluster as per global analysis, was not prominent (only 3.5% of samples) in our study population (**Fig: 7**). In addition to 19B, we found another clade emerging out of 19A, encompassing 8.8% of our sample population. This clade is defined by a set of four mutations; C28311T, C6312A, C13730T and C23929T. This clade was also identified in another Indian study^12^ where they observed around 29% of their study population to be in this cluster and named it as Clade I/A3i (defined by a combination of same four variants). This study used viral genomes isolated from Telangana along with other available sequences to infer the phylogeny. The observed differences in the frequency of the sub-populations in these two studies in Clade I/A3i is due to the altered demographics of the virus sequences.

**Fig: 7.**
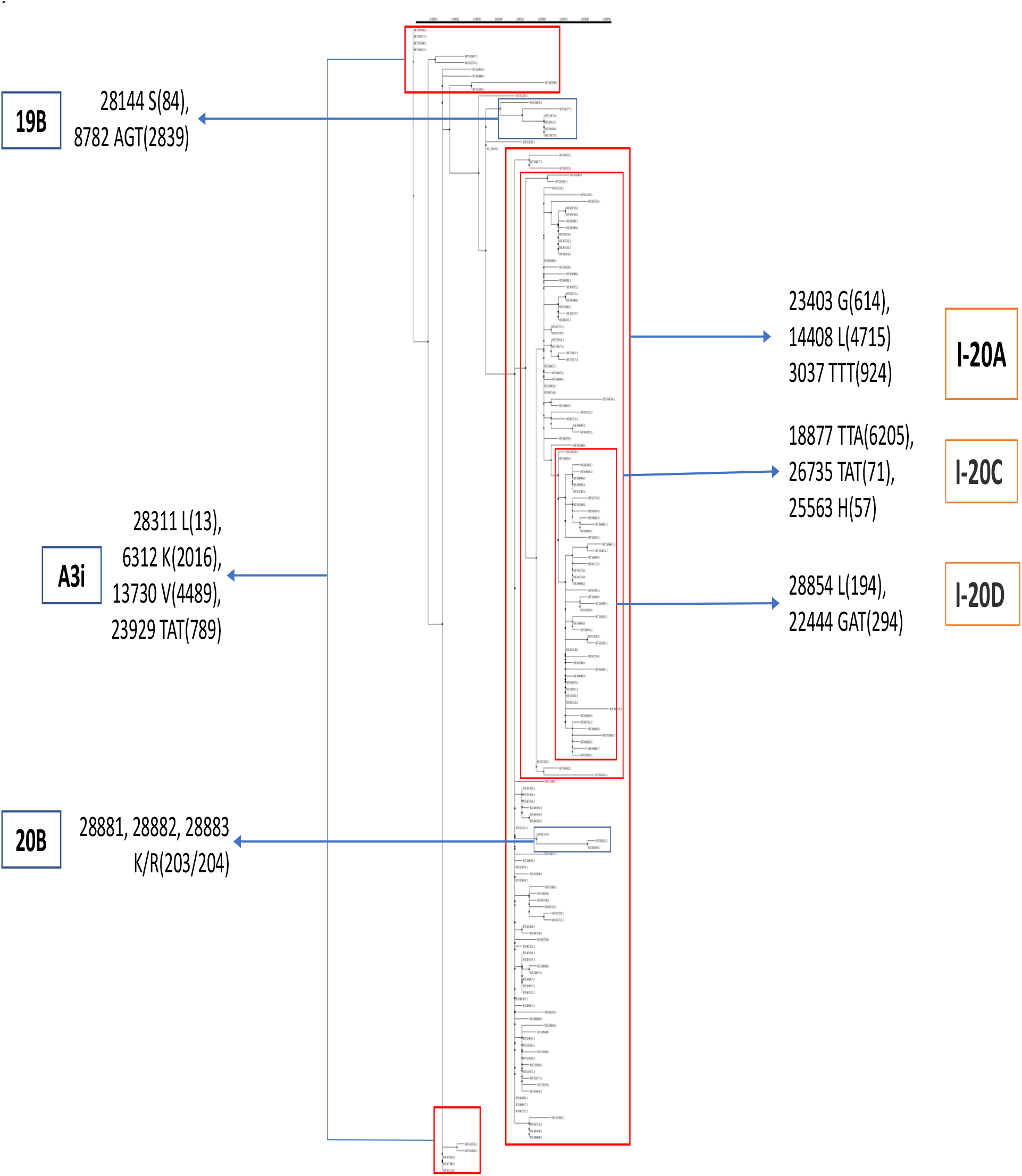
Phylogenetic analysis (100 bootstrap replications) of 174 SARS-CoV-2 genome sequences (final set). Red boxes highlight the cluster of sequences sharing linked SNPs. Blue Boxes display the global clades, Orange Boxes display clades defined in Indian population.

Most of the sequences (~ 86%) used in our study population fall in the major clade 20A (**Fig: 7**). Nextstrain^3^ defined clade 20A by two mutations, C14408T and A23403G. We found an additional mutation (C3037T) redefining the clade (I-20A, i.e. India-20A) for Indian population (high D’ and R^2^ values, Table: 2). Also, C241T mutation in the 5’UTR (not shown here) is tightly linked with these variants of clade I-20A. Similar to 19B, very few sequences (1.7%) represent clade 20B in our study population.

Clade 20C emerged from 20A and was defined by Nextstrain with two mutations, C1059T and G25563T. Surprisingly, in our study population the linkage expected between these two mutations was not found. On the contrary, G25563T was found to be strongly linked with mutations C18877T and C26735T. Since this linkage was different from what was observed for 20C, we renamed this clade as I-20C (India-20C) for Indian population.

In addition to the above clades we identified a novel clade (I-20D) emerging from I-20C, representing around 28% of the sequences in the population studied. This clade was defined by a combination of C22444T and C28854T variants (**Fig: 7**). C2836T was also found to be linked with C22444T and C28854T (**Table: 2**, **Fig: 5**). Since the allele frequency of C2836T was lower than C22444T and C28854T (Table: 1) in the final sequence set, it was not included in the variant combination defining Clade I-20D.

The analysis was further extended to global population to check the spread of the Clade I-20D across the world. SARS-CoV-2 genome sequences from various countries/continents were retrieved from NCBI and the presence of variant combination (C22444T and C28854T) was tested at respective positions of each sequence. We found this clade to be absent in Rest of Asia (Excluding India), Africa and South America whereas North America, Europe and Oceania had very low frequency (0.3, 0.3 and 0.8% respectively) of the variant combination (**Table: 3**).

**Table: 3.** Global distribution of Clade I-20D. N = Population size

| Country/Continent | Percentage of sequences showing linkage between C22444T and C28854T (update: till 05-06-2020) |
| --- | --- |
| India | 28.3% (N=173) |
| Rest of Asia | 0% (N=174) |
| North America | 0.3% (N=2570) |
| Africa | 0% (N=11) |
| Europe | 0.3% (N=288) |
| Oceania | 0.8% (N=365) |
| South America | 0% (N=6) |

### Functional evaluation of SNPs in Clade I-20D

One of the two mutations defining the clade I-20D is synonymous (C22444T), while the other one (C28854T) is non-synonymous therefore can cause a change in the amino acid sequence. The C28854T variant was analysed for potential functional consequences using the Protein Variant Effect Analyser (PROVEAN)^20^ and PolyPhen-2^21^. This non-synonymous mutation occurs in the nucleocapsid phosphoprotein, N, resulting in a change of Serine (S) to Leucine (L) at 194 position of polypeptide. The change observed is in the serine rich region (181 – 213)^15^ of the protein. Since N is a phosphoprotein and serine being a potential site for phosphorylation, change in serine to leucine can have functional consequences. Both PROVEAN and PolyPhen-2 analyses suggest that the effect of this mutation was deleterious in nature (**Table: 4**). However, further experimental studies are needed to determine the true effect of this mutation.

**Table: 4.** Characterizing the effect of the non-synonymous mutation defining Clade I-20D

| Software | Score | Prediction |
| --- | --- | --- |
| PolyPhen-2 | 0.994 (sensitivity: 0.69; specificity: 0.97) | PROBABLY DAMAGING |
| PROVEAN | -4.272 (cutoff = -2.5) | Deleterious |

## Discussion

In this report, we investigated the pattern of molecular evolution of 173 SARS-CoV-2 genomes isolated from Indian population. On the basis of linkages between sixteen recurrent SNPs (each found in more than 5% sequences), sequences were clustered and subsequently clades were defined. In this study population, we found two unique clades, that were not among the major global clades as defined by Nextstrain. One of the two clades that we identified was also reported by Banu *et at.*, wherein the cluster was named, Clade I/A3i^12^. The identification of the cluster was defined by a combination of four variants, C28311T, C6312A, C13730T and C23929T. Similar to their observation, we found another linked SNP along with these variants but not all of the sequences in the cluster shared this variant, G11083T (D’ and R^2 values with all other four variants were less than 0.78 and 0.49, respectively), therefore excluded from the clade identifier set. The above referred study included a large sequence population from Telangana with the geographical distribution of this cluster was found to be prevalent in Telangana, Tamil Nadu, Maharashtra and Delhi. The major contributors to our study population were the SARS-CoV-2 sequences originated mostly from Gujarat. Since this clade (I-A3i) may not be predominant in Gujarat, the number of sequences belonging to the clade was very less (8.8% of the total sequences).

In our analysis, we found another unique clade specific to the Indian population (absent in other countries/continents). This clade was predominantly present in the study population (28% of the total sequences) and was defined by two tightly linked SNPs, C22444T and C28854T. We named this cluster, I-20D (India-20D), since it is derived from the parent clade I-20C (India-20C). The global clade, 20C was redefined for our study population as the combination of two variants C1059T and G25563T, defining it, was missing in the sequences we analysed. Instead, G25563T was found to be strongly linked with C18877T and C26735T and hence we renamed this clade as I-20C. Similarly, clade 20A was renamed I-20A as C14408T and A23403G was tightly linked with C3037T. Therefore, the SNPs that determine the clades in global population (Nextstrain) have shown an altered occurrence patterns in Indian population.

Recently, a study demonstrated increased infectivity of SARS-CoV-2 virus when D (Aspartic Acid) at 614 position of surface glycoprotein converted into G (Glycine)^26^. The D614G (A23403G) is one of the mutation identifiers of 20A global clade, which was found to occur in 86% of our study population. Whether this mutation has resulted in a higher infectivity in Indian population needs to be investigated further. The C to T conversion at 28854 position in the N gene of viral genome in the clade I-20D (S194L) may alter the function of the Nucleocapsid Phosphoprotein. This change was predicted to be deleterious by two programmes^21,20^. More experimental studies are needed to confirm the role of this mutation in N-gene.

This study brings out the presence of a novel cluster of mutations (I-20D) forming a distinct clade, exclusively found amongst Indian SARS-CoV-2 genomes. We also found alterations in the mutation composition among viral clusters 20A and 20C. The consequence of these variations on the fitness of the virus and hence disease aetiology remains unclear. Instead, a population genetic analysis, leading to the knowledge of polymorphisms can be better utilized for drug and vaccine development.

## Acknowledgements

We thank H. S. Misra, A. Saini, S. Uppal and R.Chitella (Molecular Biology Division, BARC) and R.Shashidhar (Food Technology Division, BARC) for the critical reading of the manuscript and the suggestions.

## References

1. Bogoch, I. I. et al. Pneumonia of unknown aetiology in Wuhan, China: potential for international spread via commercial air travel. J. Travel Med. 27, (2020).

2. Coronavirus. https://www.who.int/emergencies/diseases/novel-coronavirus-2019.

3. nextstrain/ncov. GitHub https://github.com/nextstrain/ncov.

4. Gaunt, E. R., Hardie, A., Claas, E. C. J., Simmonds, P. & Templeton, K. E. Epidemiology and Clinical Presentations of the Four Human Coronaviruses 229E, HKU1, NL63, and OC43 Detected over 3 Years Using a Novel Multiplex Real-Time PCR Method. J. Clin. Microbiol. 48, 2940–2947 (2010).

5. Cui, J., Li, F. & Shi, Z.-L. Origin and evolution of pathogenic coronaviruses. Nat. Rev. Microbiol. 17, 181–192 (2019).

6. Andersen, K. G., Rambaut, A., Lipkin, W. I., Holmes, E. C. & Garry, R. F. The proximal origin of SARS-CoV-2. Nat. Med. 26, 450–452 (2020).

7. Zhou, P. et al. A pneumonia outbreak associated with a new coronavirus of probable bat origin. Nature 579, 270–273 (2020).

8. Liu, P. et al. Are pangolins the intermediate host of the 2019 novel coronavirus (SARS-CoV-2)? PLOS Pathog. 16, e1008421 (2020).

9. Li, X. et al. Emergence of SARS-CoV-2 through recombination and strong purifying selection. Sci. Adv. eabb9153 (2020) doi:10.1126/sciadv.abb9153.

10. Tang, X. et al. On the origin and continuing evolution of SARS-CoV-2. Natl. Sci. Rev. 7, 1012–1023 (2020).

11. Laamarti, M. et al. Large scale genomic analysis of 3067 SARS-CoV-2 genomes reveals a clonal geo-distribution and a rich genetic variations of hotspots mutations. http://biorxiv.org/lookup/doi/10.1101/2020.05.03.074567 (2020) doi:10.1101/2020.05.03.074567.

12. Banu, S. et al. A distinct phylogenetic cluster of Indian SARS-CoV-2 isolates. bioRxiv 2020.05.31.126136 (2020) doi:10.1101/2020.05.31.126136.

13. Database resources of the National Center for Biotechnology Information. Nucleic Acids Res. 44, D7–D19 (2016).

14. Kumar, S., Stecher, G., Li, M., Knyaz, C. & Tamura, K. MEGA X: Molecular Evolutionary Genetics Analysis across Computing Platforms. Mol. Biol. Evol. 35, 1547–1549 (2018).

15. Lu, S. et al. CDD/SPARCLE: the conserved domain database in 2020. Nucleic Acids Res. 48, D265–D268 (2020).

16. Bradbury, P. J. et al. TASSEL: software for association mapping of complex traits in diverse samples. Bioinformatics 23, 2633–2635 (2007).

17. Metsalu, T. & Vilo, J. ClustVis: a web tool for visualizing clustering of multivariate data using Principal Component Analysis and heatmap. Nucleic Acids Res. 43, W566–W570 (2015).

18. Guindon, S. et al. New Algorithms and Methods to Estimate Maximum-Likelihood Phylogenies: Assessing the Performance of PhyML 3.0. Syst. Biol. 59, 307–321 (2010).

19. Lefort, V., Longueville, J.-E. & Gascuel, O. SMS: Smart Model Selection in PhyML. Mol. Biol. Evol. 34, 2422–2424 (2017).

20. Choi, Y., Sims, G. E., Murphy, S., Miller, J. R. & Chan, A. P. Predicting the Functional Effect of Amino Acid Substitutions and Indels. PLOS ONE 7, e46688 (2012).

21. Adzhubei, I. A. et al. A method and server for predicting damaging missense mutations. Nat. Methods 7, 248–249 (2010).

22. Kryazhimskiy, S. & Plotkin, J. B. The Population Genetics of dN/dS. PLoS Genet. 4, (2008).

23. Stumpf, M. P. Haplotype diversity and SNP frequency dependence in the description of genetic variation. Eur. J. Hum. Genet. 12, 469–477 (2004).

24. VanLiere, J. M. & Rosenberg, N. A. Mathematical properties of the r2 measure of linkage disequilibrium. Theor. Popul. Biol. 74, 130–137 (2008).

25. Hadfield, J. et al. Nextstrain: real-time tracking of pathogen evolution. Bioinformatics 34, 4121–4123 (2018).

26. L, Z. et al. The D614G mutation in the SARS-CoV-2 spike protein reduces S1 shedding and increases infectivity. (2020) doi:10.1101/2020.06.12.148726.

